# Exploratory extracellular vesicle-bound miRNA profiling to identify candidate biomarkers of chronic alcohol drinking in non-human primates

**DOI:** 10.1101/2021.07.28.454223

**Authors:** Sloan A. Lewis, Brianna Doratt, Suhas Sureshchandra, Tianyu Pan, Steven W. Gonzalez, Weining Shen, Kathleen A. Grant, Ilhem Messaoudi

## Abstract

**Background:** Long-term alcohol drinking is associated with numerous health complications including susceptibility to infection, cancer, and organ damage. However, due to the complex nature of human drinking behavior, it is challenging to determine whether alcohol use should be considered a risk factor during diagnosis and treatment. We lack reliable biomarkers of alcohol use that could be used to determine drinking behavior prior to signs of overt organ damage. Recently, extracellular vesicle-bound microRNA (EV-miRNA) have been discovered to be consistent biomarkers of conditions including cancer and liver disease.

**Methods:** In this study, we profiled the plasma EV-miRNA content by miRNA-Seq from 80 non-human primates after 12 months of voluntary ethanol drinking.

**Results:** We identified a list of up- and downregulated EV-miRNA candidate biomarkers of both heavy drinking as well as those positively correlated with ethanol dose. We further overexpressed these candidate miRNA in control primary peripheral immune cells to assess potential functional mechanisms of these EV-miRNA. We identified that overexpression of miR-155, miR-154, miR-34c, miR-450a, and miR-204 led to increased inflammatory TNFα or IL-6 production in PBMC after stimulation.

**Conclusion:** This exploratory study identified several EV-miRNA that could serve as biomarkers of long-term alcohol drinking as well as provided a mechanism for alcohol-induced peripheral inflammation.

## INTRODUCTION

Alcohol consumption is widespread in the United States with 60% of the population aged 18 and older reporting alcohol use in the last month(1). Chronic alcohol consumption has been associated with numerous adverse health outcomes, including severe organ damage (2) and cancers (3, 4). Heavy alcohol consumption is also correlated with poor vaccine response(5) and increased susceptibility to many bacterial (6-8) and viral pathogens(9, 10) indicating dysregulation of the immune system. However, the molecular mechanisms underlying immune dysfunction with chronic drinking are difficult to study because human alcohol consumption is unpredictable and challenging to monitor. Currently, blood ethanol content (BEC), aspartate aminotransferase (AST), alanine amino transferase (ALT), gamma-glutamyltransferase (GGT), carbohydrate-deficient transferrin (CDT)(11), phosphatidylethanol (PEth) (12), and mean corpuscular volume (MCV), are used as clinical indicators of alcohol use(11, 13). However, these biomarkers are limited in their sensitivity and specificity to alcohol consumption. For example, elevated AST, CDT and MCV levels were observed in only 56%, 69%, and 47%, respectively, of hospitalized patients who disclosed participation in heavy alcohol consumption(14). This creates a need for new measurable, consistent biomarkers of alcohol consumption. Recent studies have highlighted the utility of extracellular vesicle bound micro RNA (EV-miRNA) as biomarkers of different disease states (15). miRNA are stable, regulatory small non-coding RNA, typically 22 nucleotides long, and are responsible for the targeting and silencing of messenger RNA (mRNA) by transcriptional blockade for degradation(16). These extracellular miRNA are able to exist independently in biofluids or packaged in extracellular vesicles, where they are stable over time(17, 18). Extracellular vesicles such as exosomes and microvesicles are found in all biofluids including serum and plasma and facilitate cell to cell communication by horizontal transfer of vesicle content, including proteins, mRNA, and many types of non-coding RNA(19).

Ethanol exposure may lead to increased production of exosomes as well as altered expression of EV-miRNA content (20). For example, miR-122, a liver-specific miRNA, is upregulated in exosomes isolated from human sera immediately following the consumption of 2 ml vodka 40% v/v ethanol/kg body weight and in mice models of binge drinking (5 g/kg 50% v/v ethanol diluted in water via oral gavage), and chronic drinking (a diet consisting of 36% of calories coming from ethanol for 5 weeks) (20). Also, increased abundance of miR-122 results in monocyte hypersensitivity to lipopolysaccharides (LPS) (20). Treatment of THP-1 cells in vitro with alcohol increases miR-27a abundance in EVs, which in turn induces M2 polarization in naïve THP-1 cells (21). Additionally, studies have reported miR-155, miR-122, and miR-34a as a potential biomarkers of alcohol-induced liver disease (20, 22, 23). However, the identification of EV-miRNA that could be used as reliable biomarkers of chronic ethanol consumption in the absence of overt organ damage is challenging when using strictly clinical samples, rodent models, or treated cell lines. Clinical studies are limited by inaccurate self-reporting of alcohol intake and medical histories, concurrent drug use, and environmental factors. Data from rodent model studies is confounded by the stress caused by either forced feeding with surgically placed gavage tubes, access to high concentration ethanol only diets, and only short periods of ethanol exposure.

In the presented study, we utilized a rhesus macaque model of voluntary ethanol self-administration to identify potential extracellular vesicle bound miRNA (EV-miRNA) biomarkers of alcohol drinking. This model allows for elucidation of the impact of chronic alcohol consumption in an outbred species while avoiding confounding factors such as smoking or drug use. Using this model, we have reported large transcriptomic changes in peripheral blood mononuclear cells (PBMC) and tissue resident macrophages with alcohol consumption (24, 25). These studies revealed altered expression of genes associated with host defense, inflammation, and wound repair (26, 27). This transcriptional dysregulation could be partially explained by altered miRNA expression that we have observed in PBMC from the same animals (25-27), however, global alterations in circulating and extracellular vesicle bound miRNA have not been explored. Therefore, we examined the influence of alcohol drinking on EV-miRNA expression using samples obtained from 80 rhesus macaques after 12 months of open access to an alcohol solution and controls. Our analysis revealed that chronic ethanol consumption results in altered EV-miRNA cargo. We were able to identify several EV-miRNA that were up-or down-regulated with alcohol drinking and that regulated genes with roles in myeloid cell activation, angiogenesis, and regulation of gene expression. Eight EV-miRNA were selected for additional functional studies. Over-expression of candidates miR-155, miR-154, miR-34c, miR-450a, and miR-204, which are upregulated in EV with alcohol drinking, resulted in a heightened inflammatory response in PBMC after stimulation. These findings both provide candidates that can be further validated as biomarkers of chronic alcohol drinking as well as provide a potential mechanism for increased inflammation associated with alcohol use.

## RESULTS

### Extracellular vesicle associated miRNA (EV-miRNA) expression changes with heavy alcohol drinking

To identify candidate biomarkers of alcohol drinking, we profiled miRNA from circulating extracellular vesicles (EV-miRNA) isolated from macaque plasma after 12 months of voluntary alcohol drinking (**Figure 1A**). Total RNA was extracted from the extracellular vesicles and miRNA libraries were constructed for 22 control, 33 Chronic Moderate Drinking (CMD), and 25 Chronic Heavy Drinking (CHD) animals (**Supp. Table 1**). Initial analysis of the sequencing data revealed significant differences between male and female animals, therefore separate analyses were carried out for each. We first sought to discover expression changes in EV-miRNA with chronic heavy alcohol drinking (CHD; >3g EtOH/kg bodyweight/day). Differential analysis of miRNA-Seq data from CHD and control animals revealed a number of differentially expressed miRNA (DEmiRNA) in both males and females (**Figure 1B,C**). Only one upregulated (miR-154) and one downregulated (miR-1224) DEmiRNA were found to be in common between males and females by this analysis (**Figure 1D**). We next identified validated gene targets of the human homologs of these DEmiRNA using miRTarBase and performed functional enrichment on those genes using Metascape (28) (**Figure 1E and Supp. Figure 1A**,**B**). Although the numbers of validated gene targets varied greatly between EV-miRNA (**Supp. Figure 1A**,**B**), we were able to identify significantly enriched pathways from each target gene list. The targets of several upregulated EV-miRNA enriched to gene ontology (GO) terms associated with blood vessel development such as *angiogenesis* and *vasculature development* (e.g., miR-124a, 130b, 141 and 34c). Several EV-miRNA targeted genes involved in the *response to reactive oxygen species* (e.g. miR-124a ,433, 493 and 34c). Others targeted genes that play a role in *myeloid leukocyte activation* (e.g. miR-124a, 130b, 154, 208, 544) (**Figure 1E**). We also performed functional enrichment on the targets of the downregulated EV-miRNA to assess processes that may be upregulated with CHD. We found the targets of these downregulated EV-miRNA mapped to Go terms associated with signaling (e.g., “regulation of protein kinase activity”), cell cycle (e.g., “mitotic DNA damage checkpoint”), and tissue/epithelial homeostasis (e.g., “tissue homeostasis” and “response to hypoxia”) (**Supp. Figure 1C**).

**Figure 1:**
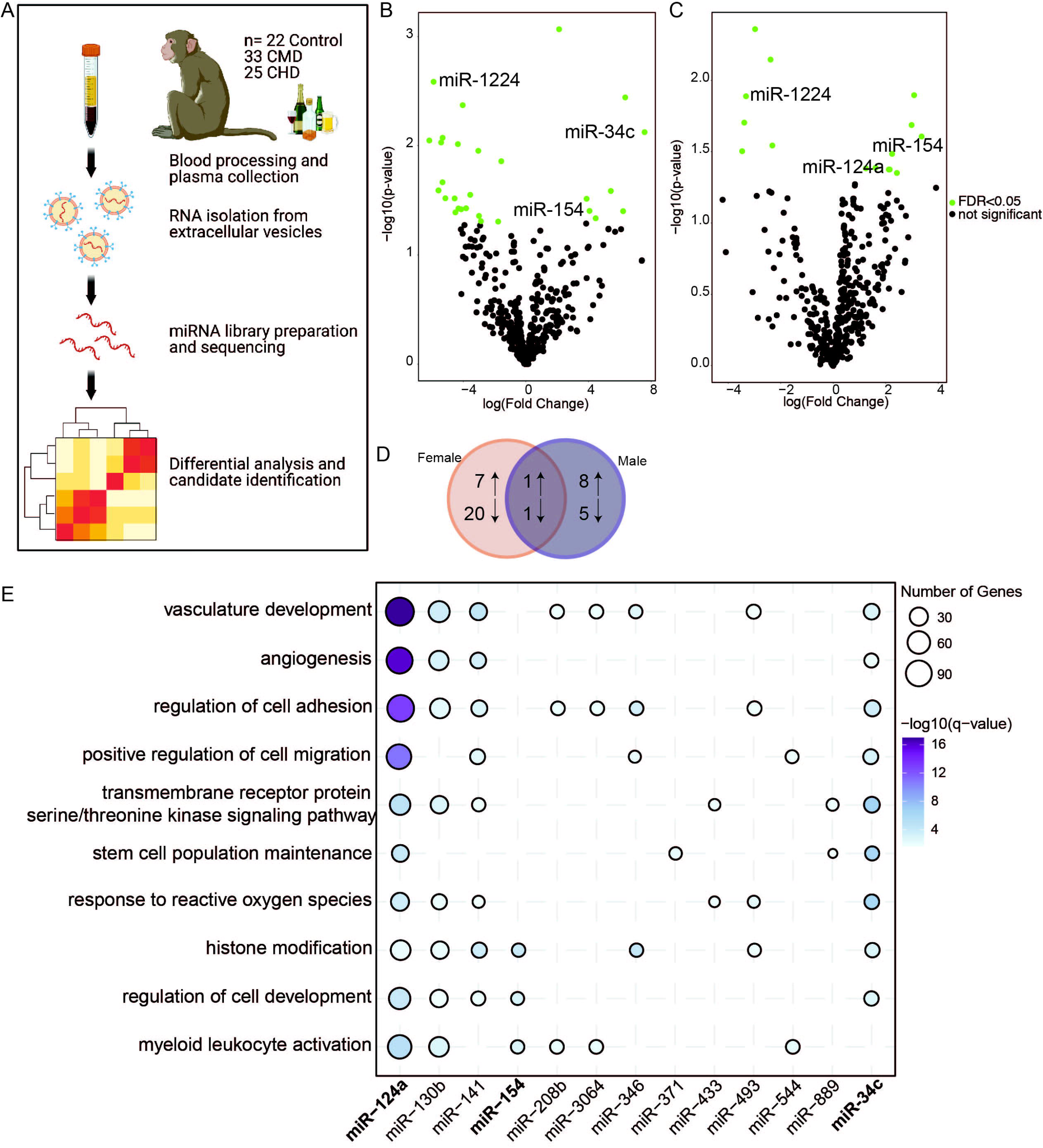
Extracellular vesicle associated miRNA (EV-miRNA) expression changes with heavy alcohol drinking. A) Experimental Design for the study. B,C) Volcano plots showing significant (FDR≤0.05) differentially expression miRNA with heavy drinking in female (B) and male (C) macaques. D) Venn diagram comparing DEmiRNA between male and female macaques with heavy drinking. E) Bubble plot depicting GO Biological processes associated with the validated target genes of the indicated upregulated EV-miRNA with heavy drinking. The size of the bubble represents the number of target genes, and the color represents the -log_10_(q-value) significance of enrichment.

### Dose-dependent changes in extracellular vesicle associated miRNA expression

We next identified EV-bound miRNA whose expression was significantly correlated with dose of ethanol (g/kg/day). Similar trends in male and female animals were noted; therefore, data from both sexes were pooled for this analysis (**Figure 2A**). We identified 12 miRNAs whose expression was positively correlated with ethanol dose (**Figure 2A**). The expression of one EV-miRNA, miR-155, was only positively correlated with ethanol consumption in female animals (**Figure 2B**). We used miRTarBase to identify validated gene targets of each of these miRNAs (**Supp. Figure 2A**) and carried functional enrichment to understand the biological processes impacted by these DEmiRNA (**Figure 2C**). The targets of several EV-miRNA enriched to gene ontology (GO) terms associated with blood vessel development such as *Blood vessel morphogenesis* and *vasculature development* (e.g., miR-335, 204, 155) (**Figure 2B**). A few EV-miRNA targeted genes regulate tissue repair as indicated by enrichment to GO terms *response to wounding, epithelial cell proliferation* and *cellular response to growth factor* stimulus (e.g., miR-211, 335, 204, 96) (**Figure 2B**). EV-miRNA also targeted genes mapping to *myeloid cell differentiation* (e.g., miR-335, 579, 454, 197, 155).

**Figure 2:**
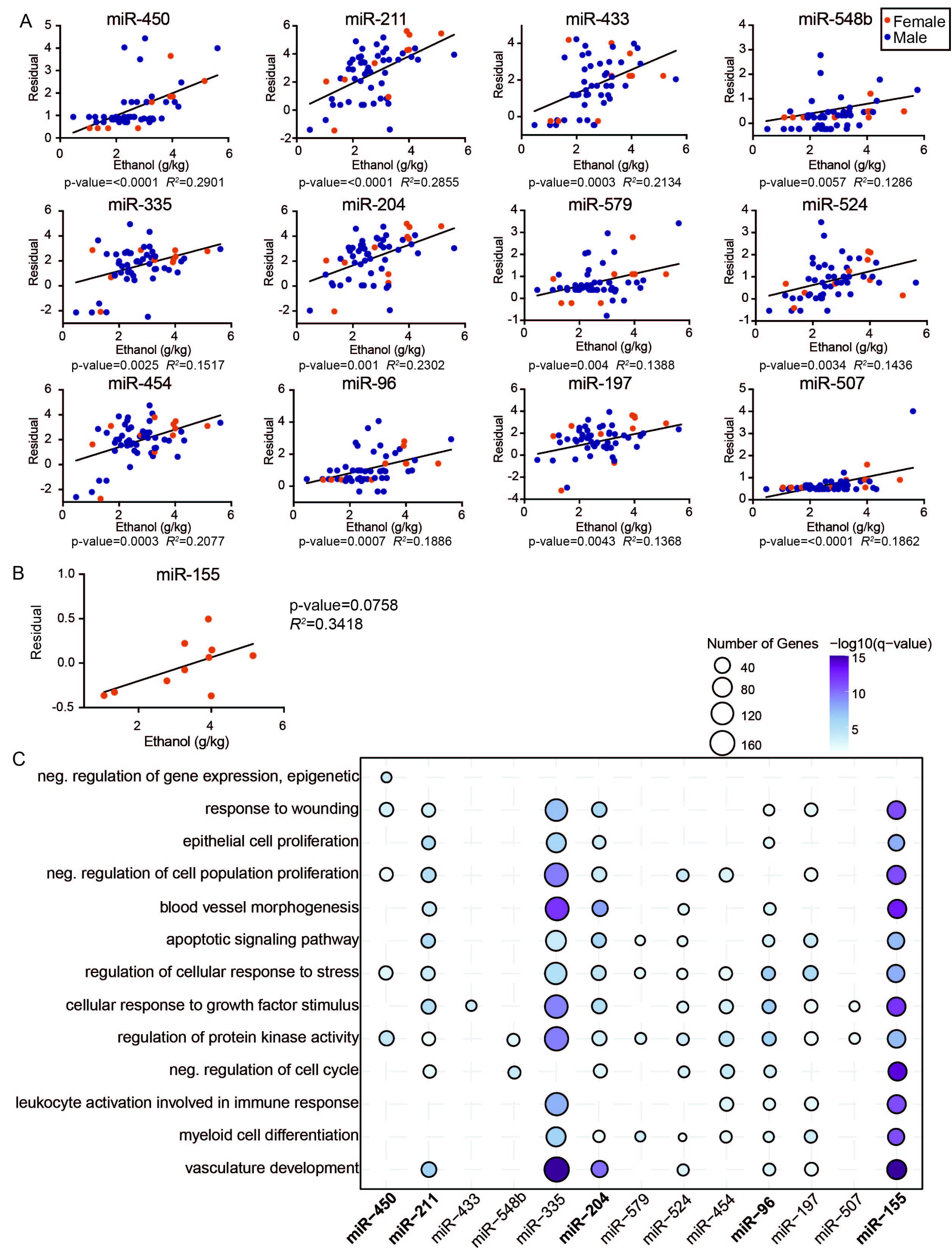
Dose-dependent changes in extracellular vesicle associated miRNA expression. A) Scatter plots of partial residual expression values for selected miRNA and ethanol dose (g EtOH/ kg body weight/ day). B) Scatter plots of partial residual expression values for miR-155 and ethanol dose (g EtOH/ kg body weight/ day) in female animals. C) Bubble plot depicting GO Biological processes associated with the validated target genes of the indicated positively correlated EV-miRNA with ethanol dose. The size of the bubble represents the number of target genes, and the color represents the -log_10_(q-value) significance of enrichment.

### Upregulated EV-miRNA associated with heightened inflammatory response in PBMC

The analyses described above identified a total of 28 upregulated and 27 downregulated potential candidate EV-miRNA to predict alcohol consumption (**Supp. Table 2**). We next asked whether these circulating EV-miRNA impact the gene expression of PBMC. We compared the list of down-and up-regulated differentially expressed genes (DEG) reported in our previous studies of PBMC (25) to that of the validated target genes of the up-and down-regulated EV-miRNA respectively (**Figure 3A,B**). This analysis showed that 334 of the validated targets of upregulated EV-miRNA were previously identified as downregulated DEG in PBMC from alcohol consuming animals (25) (**Figure 3A**). Functional enrichment of these common DEG enriched to GO terms associated with regulation of TGFβ receptor signaling and chromatin modification (**Figure 3C**). The 85 validated targets of downregulated EV-miRNA that were shared with DEG previously reported (25) to be upregulated with chronic heavy drinking enriched to apoptotic/DNA damage and coagulation related processes (**Figure 3B,C**). To evaluate whether these EV-miRNA were derived from PBMC themselves or other tissues, we compared the up-and downregulated EV-miRNA to differentially expressed cellular miRNA from male and female PBMC with alcohol drinking (25) (**Supp. Figure 2B**). We identified only 2 upregulated and 1 downregulated miRNA in common, suggesting that the increased or decreased EV-miRNA in circulation are most likely coming from tissues other than PBMC (**Supp. Figure 2B**).

**Figure 3:**
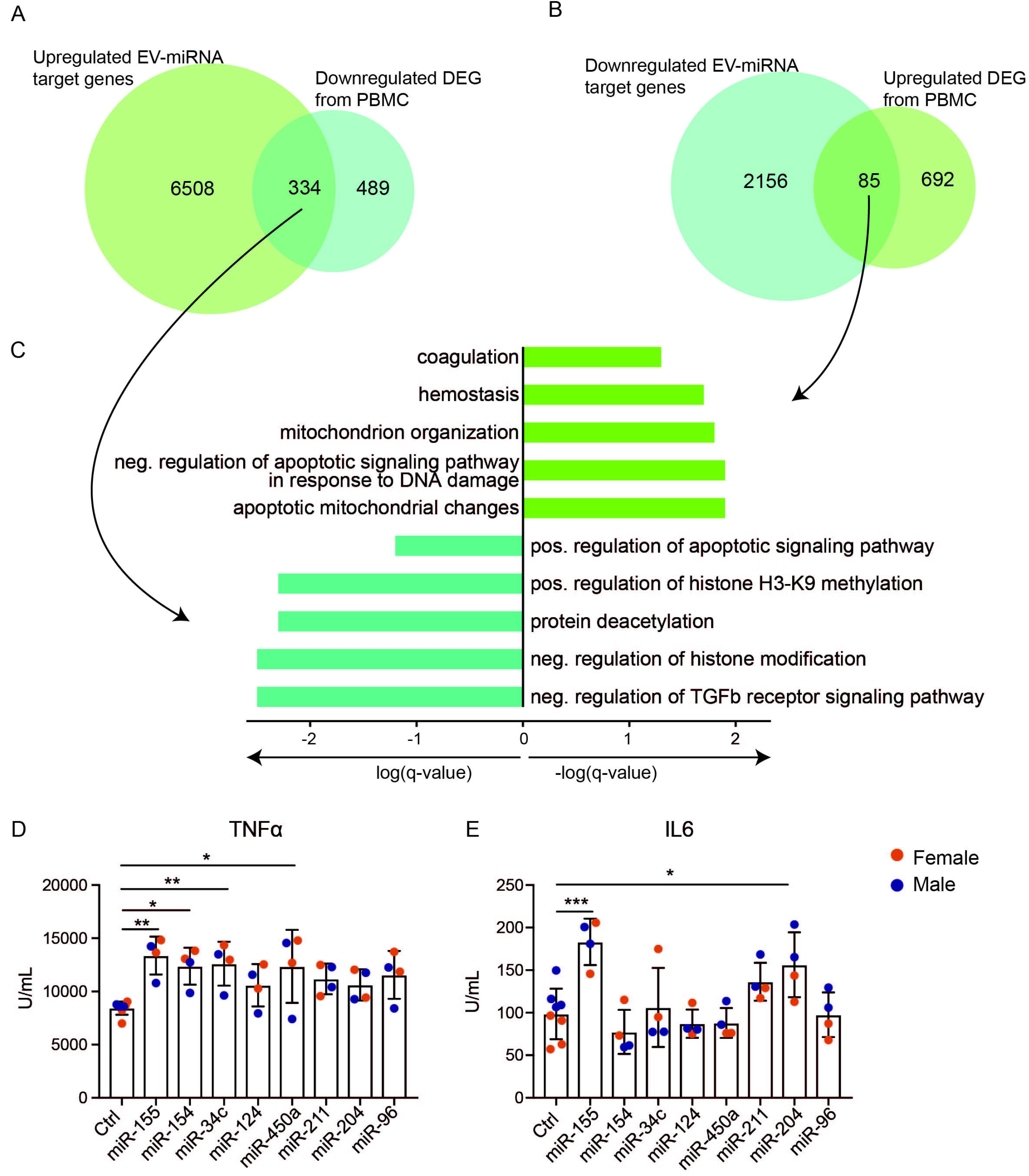
Upregulated EV-miRNA associated with heightened inflammatory response in PBMC. A) Venn diagram comparison of target genes from upregulated EV-miRNA with downregulated DEG from PBMC (25). B) Venn diagram comparison of target genes from downregulated EV-miRNA with upregulated DEG from PBMC. C) GO Biological process enrichment of genes in common from (A) and (B) where the X-axis is log_10_(q-value) or -log_10_(q-value), respectively. D,E) PBMC from control macaques were transfected with miRNA mimics and further stimulated with PMA/ionomycin. The production of TNFα (D) and IL-6 (E) was measured from supernatants by ELISA. P-value significance is denoted by *=<0.05, **=<0.01, ***=<0.001.

A more focused list of 8 EV-miRNA was selected based on the validated target genes of these EV-miRNA as well as literature searches on the EV-miRNA themselves (**Supp. Table 3**). To assess the functional implications of these 8 EV-miRNA on circulating immune cells, we transfected PBMC obtained from ethanol naïve male and female macaques (n=2/sex) with their mimics (**Supp. Figure 2C**). We then stimulated the cells with PMA/ionomycin and assessed production of canonical pro-inflammatory cytokines TNFα and IL-6. PBMC transfected with miR-155, generated a heightened IL-6/TNFα response, whereas PBMC transfected with miR-154, miR-34c, or miR450a mimics produced increased amounts of TNFα (**Figure 3D,E**). Cells transfected with miR-204 produced increased levels of IL-6 only (**Figure 3E**). These results indicate that increased levels of these extracellular vesicle bound miRNA in circulation with chronic drinking could lead to heightened inflammatory mediator production in response to stimulation.

## DISCUSSION

Long term alcohol drinking is a complicating factor for numerous health conditions including infections, cancer, and organ damage (1, 29). However, a lack of unbiased methods for measuring alcohol drinking over time makes assessing the impact of chronic ethanol consumption challenging to assess. Our first goal of this study was to utilize a rhesus macaque model of voluntary ethanol self-administration to discover candidate extracellular vesicle bound miRNA (EV-miRNA) biomarkers of chronic alcohol drinking. We explored potential candidates of heavy drinking as well as candidates that correlated with average dose of ethanol. In this model, chronic heavy drinking is defined by The Monkey Alcohol Tissue Research Resource (MATRR) as >3 grams of ethanol per kilogram body weight per day. As one standard drink is 14 grams of ethanol, this equates to more than 12 drinks per day in an average 60kg human. Due to known differences in alcohol metabolism and long-term effects between males and females, we first examined the EV-miRNA profiles of these heavy drinking macaques after 12 months of this behavior separated by sex. This allowed us to identify sex-dependent and independent EV-miRNA. A total of 16 EV-miRNA were upregulated while 26 were downregulated with chronic heavy drinking with only 2 EV-miRNA shared between the two groups (miR-154 was upregulated and miR-1224 was downregulated). Validated targets of the upregulated EV-miRNA regulated expression of genes involved in pathways associated with blood vessel development as well as cell migration and adhesion. As these EV-miRNA are in circulation, their functional implications could be widespread throughout the body.

The macaque model allows for a variety of drinking patterns and behaviors and therefore we were able to profile the plasma EV-miRNA of animals who consumed alcohol more moderately over the yearlong drinking period (<3g EtOH/ kg body weight/ day). We further correlated miRNA expression with dose of ethanol to identify 13 miRNA positively associated with alcohol dose. Interestingly, these miRNA, like those associated with heavy drinking, also targeted genes involved in blood vessel development and wounding processes. Accordingly, both moderate and heavy alcohol drinking have been linked to hypertension, cardiovascular disease and heart failure in men and women (30-32) and EV-miRNA have been implicated in cardiovascular maintenance and development of disease (33). Whether these alcohol-induced EV-miRNA are directly targeting these processes requires further investigation.

As alcohol and its metabolites circulate through the body and act on both peripheral and tissue-resident cell populations, the cellular origins of these EV-miRNA are difficult to parse. To account for peripheral immune cells as a source, we compared the differential EV-miRNA to differential miRNA identified in PBMC with alcohol drinking (25). No significant overlap was seen between EV-miRNA and PBMC miRNA, strongly suggesting that the source of the EV-miRNA is likely to be non-immune tissues. Previous studies have identified the pancreas, liver, intestine, brain, and heart to have altered miRNA expression with ethanol exposure (34). Further, studies have identified liver-derived EV-bound miRNA 122 and let7f as indicators of alcohol-induced liver injury (35). While it is known that miRNA packaging in EVs is selective and can have targeted effects on recipient cells, many of the functional mechanisms and downstream impacts of these processes remain elusive (36, 37).

We and others have shown chronic alcohol drinking to alter the function of peripheral immune cells, poising them towards a hyper-inflammatory response (25, 29). We sought to determine whether the EV-miRNA that became more abundant with alcohol drinking could be modulating the function of PBMC. We first compared the validated gene targets of the EV-miRNA with the DEG identified from PBMC obtained from some of the same animals (25) and identified that the upregulated EV-miRNA targets correlated with regulation of TGFβ signaling and histone modification pathways in PBMC. We narrowed our focus to 8 potential candidate EV-miRNA that were either upregulated with heavy drinking or positively correlated with daily ethanol consumption dose and had regulated expression of genes in inflammatory pathways (miR-155, miR-154, miR-34c, miR-124, miR-450a, miR-211, miR-204, and miR-96). Overexpression of miR-155, miR-154, miR-34c, and miR-450a primary macaque PBMC led to increased TNFα production following stimulation with PMA, whereas miR-155 and miR-204 led to increased IL-6 production. The mechanisms by which these miRNAs modulate production of IL-6 and TNFα could be occurring through either a direct inhibition of NFκB regulators or via an indirect/secondary target. For instance, miR-155 has previously been shown to increase inflammatory responses through direct targeting of NFKB pathway regulators (*BCL6*) and pro-inflammatory macrophage activation (38, 39). The miR-34 family members have been linked to inflammation in the skin and liver and have been associated with non-alcoholic fatty liver disease (22, 40, 41). Increased expression of these miRNA in EV could indicate early signs of alcohol-induced liver damage prior to increased levels of AST or ALT.

In this study, we comprehensively profiled the blood plasma of macaques to find potential EV-miRNA biomarkers of long-term alcohol drinking and provide a mechanism of alcohol-associated inflammation in peripheral immune cells. Our exploratory examination led to the discovery of potential candidate biomarkers of chronic alcohol consumption in macaques. To validate these candidates as true biomarkers, a confirmatory study would need to be performed in a new cohort of animals and then further in humans. While miRNA are generally well-conserved across species, the differences between the EV-miRNA in macaques and humans has yet to be investigated.

## MATERIALS AND METHODS

### Animal studies and sample collection

These studies used samples from a macaque model of voluntary ethanol self-administration established through schedule-induced polydipsia (42-44). Briefly, in this model, rhesus macaques are introduced to a 4% w/v ethanol solution during a 90-day induction period followed by concurrent access to the 4% w/v solution and water for 22 hours/day for one year. During this time, the macaques adopt a stable drinking phenotype defined by the amount of ethanol consumed per day and the pattern of ethanol consumption (g/kg/day) (42). Blood samples were taken from the saphenous vein every 5-7 days at 7 hours after the onset of the 22 hours/day access to ethanol and assayed by headspace gas chromatography for blood ethanol concentrations (BECs).

For these studies, blood plasma from a total of 80 rhesus macaques (22 controls, 33 moderate, and 25 heaving ethanol drinking animals); 17 females and 71 males across 8 cohorts was utilized. Drinking classifications were defined as chronic moderate drinking (CMD) or chronic heavy drinking (CHD) based on 12 months of ethanol self-administration (tissue and drinking data obtained from the Monkey Alcohol Tissue Research Resource: www.matrr.com). These cohorts of animals (Cohorts 4, 5, 6a, 6b, 7a, 7b, 10a, and 14 on matrr.com) were described in two previous studies of peripheral immune system response to alcohol (24, 25). Blood plasma was isolated by centrifugation over histopaque (Sigma, St Louis, MO) as per manufacturer’s protocol and stored at -80C until they could be analyzed as a batch. The average daily ethanol intake for each animal is outlined in **Supp. Table 1**.

### Plasma EV Isolation and miRNA library preparation

Plasma was filtered using Viviclear Mini Centrifugal Filters (Sartorius, Göttingen, Germany), a 0.8µm porous membrane. Extracellular vesicle (EV)-bound total RNA was isolated from the vesicles using the exoRNeasy Serum/Plasma Midi Kit (Qiagen, Valencia, CA) by membrane affinity column purification per the manufacturer’s protocol. EV-miRNA were selected by specific ligation of adaptors to the 3’ hydroxyl and 5’ phosphate groups, unique to mature miRNA, using the QIAseq miRNA Library Kit (Qiagen, Valencia, CA). Briefly, with a 5 μl total RNA input, cDNA synthesis was performed using universal reverse transcription, library amplification, and library cleanup.

### Sequencing, identification of DEmiRNA, DEmiRNA targets, and enrichment

Each library was indexed using a unique barcode for multiplexing and sequenced on the NextSeq2500 or Hiseq4000 platform (Illumina, San Diego, CA) to yield single-end 100 bp sequences. Quality reports for the raw miRNA reads were generated using FASTQC. The reads were trimmed 10-50 bp. Followed by alignment to the *Macaca mulatta* genome from Ensembl using splice aware short read aligner suite Bowtie 2 (45). The transcript counts per gene were then summarized using the *summarizeOverlaps* function. Samples were required to have a minimum of 1,500,000 reads post alignment and 400 detected miRNA to ensure good co verage of the genome. Differentially expressed miRNA (DEmiRNAs) were identified using the edgeR package (46) with batch correction and were defined as those with a fold change ≥ 1.5, FDR corrected p-value ≤ 0.05. DEmiRNA from macaques were converted to their human homologs. The miRTarBase database (47) was scanned with the list of DEmiRNAs to identify validated gene targets. Functional enrichment of targeted genes was done using Metascape (28) to identify clusters of genes mapping to specific biological or molecular pathways. ggplot2 was used for enrichment visualization.

### Electroporation with miRNA mimics and stimulation of control PBMC

Total PBMC from control male and female macaques were thawed and washed four times in Optimem (ThermoFisher, Waltham, MA) to remove traces of serum. 5×10^6^ cells were resuspended in 200ul Optimem containing 2ug miRNA mimic (Dharmacon, Lafayette, CO). The Gene pulser II System (Bio-Rad Laboratories, Berkeley, CA) was used to electroporate the sample in a 0.4cm gap cuvette using 500V for 0.05 seconds. Cells were moved to a 48 well plate containing 500uL RPMI supplemented with 10% FBS, 1% L-Glutamine, and 1% Penicillin/Streptomycin and allowed to recover for 48 hours in an incubator with 5% CO2 at 37C. Cells were then stimulated for 16 h with or without PMA and Ionomycin at 37°C in an incubator with 5% CO2. Supernatants were collected and cells were resuspended in Qiazol (Qiagen, Valencia CA) for RNA extraction. Both cells and supernatants were stored at -80 °C until they could be processed as a batch.

### TNF and IL-6 ELISA

Supernatants were collected following 16 h of incubation. Samples were analyzed in duplicates using Monkey IL6 ELISA and Monkey TNFα Total ELISA kits (ThermoFisher, Waltham, MA) per the manufacturer’s instructions. Unstimulated condition measured values were subtracted from stimulated condition values to plot a corrected value for each test group and control.

### qPCR for electroporation efficiency

Cells were lysed in QIAzol Lysis reagent (Qiagen, Valencia, CA) and total RNA was extracted using the miRNeasy mini kit (Qiagen, Valencia, CA) per manufacturer’s instructions. We used TaqMan miRNA Assays (ThermoFisher, Waltham, MA) for detection of miRNA. Briefly, reverse transcription (30 min, 16 °C; 30 min, 42 °C; 5 min 85 °C) was done using 10ng RNA, TaqMan primers and High Capacity cDNA Reverse Transcription kit (ThermoFisher, Waltham, MA). qPCR was done on the StepOnePlus Real-Time PCR System (ThermoFisher, Waltham, MA) using TaqMan Universal PCR Master Mix.

### Statistical Analysis

All statistical analyses were conducted in Prism 7(GraphPad). Data sets were first tested for normality. One-way ANOVA ((=0.05) followed by Holm Sidak’s multiple comparisons tests was used to detect significance between control and test groups. Error bars for all graphs are defined as ± SEM. Multivariate linear regression analysis compared significant shifts in curve over horizontal line, with R-squared reported. Statistical significance of functional enrichment was defined using hypergeometric tests. P-values less than or equal to 0.05 were considered statistically significant. Values between 0.05 and 0.1 are reported as trending patterns.

## Author Contributions

S.A.L., K.A.G., and I.M. conceived and designed the experiments. S.A.L., B.D., and S.W.G. performed the experiments. S.A.L., B.D., and T.P. analyzed the data. S.A.L. and I.M. wrote the paper. All authors have read and approved the final draft of the manuscript.

## Acknowledgements

We are grateful to the members of the Grant laboratory for expert animal care and sample procurement. We thank Dr. Jennifer Atwood for assistance with sorting in the flow cytometry core at the Institute for Immunology, UCI. We thank Dr. Melanie Oakes from UCI Genomics and High-Throughput Facility for assistance with 10X library preparation and sequencing.

## Data availability

The datasets supporting the conclusions of this article are available on NCBI’s Sequence Read Archive (SRA# Pending).

## SUPPLEMENTAL FIGURE LEGENDS

**Supp. Figure 1:** A,B) Bar graph representing the number of validated target genes for each of the up- (A) or down-regulated (B) DEmiRNA identified with heavy drinking. C) Bubble plot depicting GO Biological processes associated with the validated target genes of the indicated downregulated EV-miRNA with heavy drinking. The size of the bubble represents the number of target genes, and the color represents the -log_10_(q-value) significance of enrichment.

**Supp. Figure 2:** A) Bar graph representing the number of validated target genes for each of the EV-miRNA positively correlated with ethanol dose. B) Venn diagrams of up-and downregulated EV-miRNA compared to up-and downregulated PBMC miRNA (25). C) Ct values for qPCR confirming the transfection of miR-155 and miR-211 into PBMC. P-value significance is denoted by ****=<0.0001.

## SUPPLEMENTAL TABLES

**Supp. Table 1**: Cohort characteristics for macaque plasma samples.

**Supp. Table 2**: Overall findings of EV-miRNA associated with alcohol drinking in macaques

**Supp. Table 3**: Literature review of candidate miRNA biomarkers of ethanol drinking

