## Supplementary figures and images for "Exploratory extracellular vesicle-bound miRNA profiling to identify candidate biomarkers of chronic alcohol drinking in non-human primates"

### Supp Figure 1

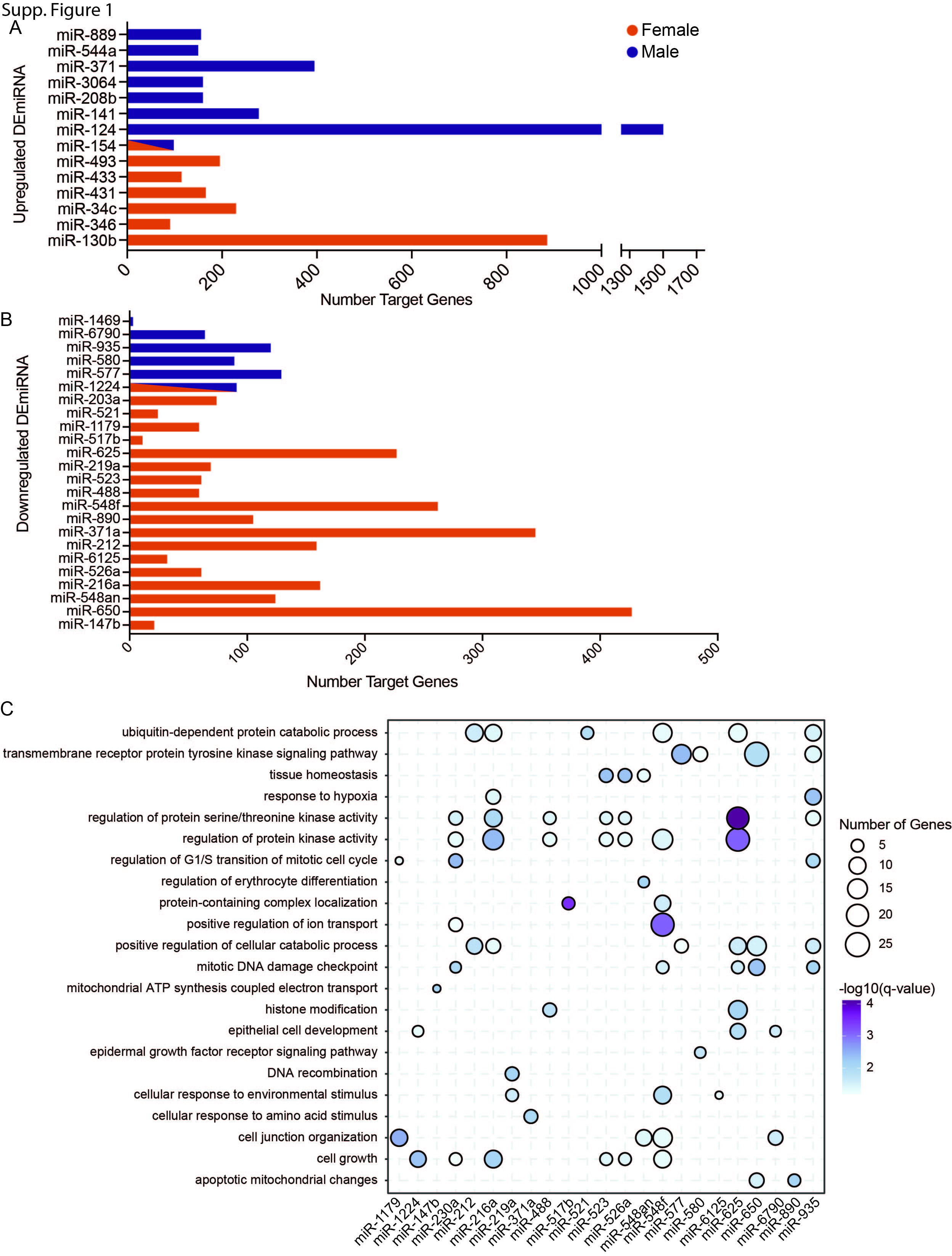

### Supp Figure 2

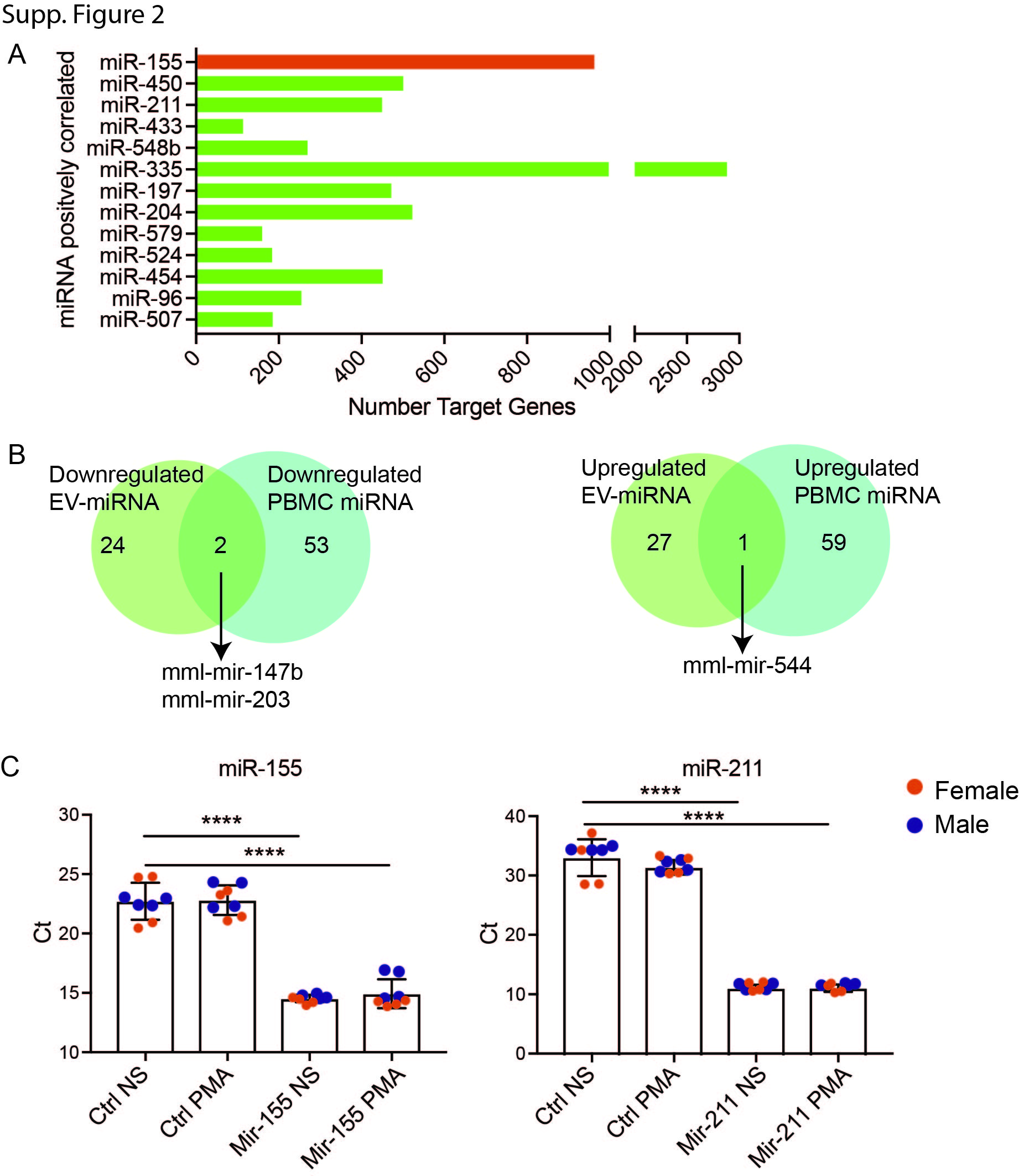
